# GABAergic malfunction underlying maternal immune activation-induced social deficits

**DOI:** 10.1101/319244

**Authors:** Kazuki Okamoto, Natsuko Hitora-Imamura, Hiroyuki Hioki, Yuji Ikegaya

## Abstract

Social deficits are one of the major symptoms of psychiatric disorders, including autism spectrum disorders (ASDs) and schizophrenia. However, the underlying mechanism remains ill-defined. Here, we focused on the anterior cingulate cortex (ACC), a brain region that is related to social behaviors, of mice that received poly(I:C)-induced maternal immune activation. Using whole-cell patch clamp recordings, we found that layer 2/3 pyramidal cells were hyperactive in acute ACC slices prepared from poly(I:C)-treated mice compared to those from saline-treated mice. The hyperexcitation was associated with a reduction in inhibitory synapse activity. Local injection of the GABA_A_ receptor enhancer clonazepam into the ACC of poly(I:C)-treated mice restored the social behaviors of the mice. These results suggest that the balanced excitability of ACC neurons is essential for social ability.

## 1. Introduction

Social deficits are observed in many developmental neuropsychiatric disorders, including autism spectrum disorder (ASD) and schizophrenia. Although the risk factors associated with these disorders are not yet strictly identified, genetic and environmental factors that are likely involved in the etiology of the disorders are often associated with synapse formation. Indeed, disruption of the excitatory and inhibitory balance (E/I balance) of synaptic activity is suggested to underlie the symptoms of psychiatric disorders (Gogolla et al., 2009). For instance, neuroimaging studies demonstrated hyperexcitability of the neocortex of ASD patients in a resting state (Agam et al., 2010, Assaf et al., 2010). Furthermore, hyperexcitation in the sensory cortex impairs sensory input processes (Gogolla et al., 2014) and induces social deficits (Yim et al., 2017). Thus, cortical hyperexcitation likely contributes to the symptoms of psychiatric disorders (Yim et al., 2017).

Previous studies suggest that a disrupted E/I balance leads to deficits in social interactions (Han et al., 2014, Yizhar et al., 2011). However, neither the responsible brain region nor the underlying synaptic mechanism is fully understood. Here, we focused on the anterior cingulate cortex (ACC), a brain region that is associated with social behaviors (Behrens et al., 2009, Singer et al., 2004). In rodents, inactivation of ACC neurons reduces empathetic behaviors (Kim et al., 2012).

In the present study, we employed a maternal immune activation (MIA) mouse model to investigate the relationships between ACC neuronal activity and social deficits. MIA is one of the major environmental factors causing psychiatric disorders in offspring (Atladottir et al., 2010, Patel et al., 2017), and social deficits are observed in MIA-model mice (Choi et al., 2016, Hsiao et al., 2013, Malkova et al., 2012). Using whole-cell patch clamp recordings, we found hyperexcitation of layer 2/3 pyramidal cells in acute ACC slices prepared from poly(I:C)-treated mice. In addition, local injection of the GABA_A_ receptor enhancer clonazepam into the ACC of poly(I:C)-treated mice restored the social behaviors of the mice. These results suggest that balanced excitation in ACC neurons is critical for social behaviors and that disruption of this balance leads to social disability.

## 2. Materials and methods

### 2.1. Animal experiment ethics

Experiments were performed with the approval of the animal experiment ethics committee at the University of Tokyo (approval no. P24-8 and P24-70) and according to the University of Tokyo guidelines for the care and use of laboratory animals. C57BL/6J mice (SLC, male) were housed in cages under standard laboratory conditions (12-h light/dark cycle, ad libitum access to food and water). All efforts were made to minimize the animals’ suffering and the number of animals used.

### 2.2. MIA model

Maternal viral infection was imitated using poly(I:C), a synthetic analogue of double-stranded RNA (Sigma) or saline (control), which was intraperitoneally injected at doses of 0.25 and 0.125 U/g on embryonic days 12.5 and 17.5, respectively (Naviaux et al., 2013). Pups were removed from their dams at 3 postnatal weeks and housed in cages of 2−4 animals until 4−6 postnatal weeks.

### 2.3. Three-chamber test

The testing apparatus measured 51 × 25 × 25 cm and was divided into three chambers (17 × 25 × 25 cm) with transparent plastic walls; the mice had free access to all the chambers through open doors. A transparent plastic cage was located in the corner of each side of the chamber. A mouse was placed in the apparatus, was allowed to explore these chambers for 10 min, and was then moved to a resting cage. Another unfamiliar mouse was placed in one of the corner plastic cages, whereas the other cage was kept vacant. In the test session, the subject mouse was returned to the testing apparatus and allowed to explore the chambers for 10 min. The testing apparatus was video-monitored. The difference between the times spent in the chamber with an unfamiliar mouse and in the vacant chamber was used as an index for social interaction.

### 2.4. Acute slice preparation

Acute slices were prepared from the ACCs of male mice (4-6 postnatal weeks). Mice were anesthetized with isoflurane and then decapitated. The brains were removed and placed in an ice-cold oxygenated solution consisting of (in mM) 222.1 sucrose, 27 NaHCO_3_, 1.4 NaH_2_PO_4_, 2.5 KCl, 1 CaCl_2_, 7 MgSO_4_, and 0.5 ascorbic acid. The brains were sliced at a thickness of 400 μm coronally for the ACC using a VT1200S vibratome (Leica). Slices were allowed to recover at room temperature for at least 1.5 h while submerged in a chamber filled with oxygenated aCSF consisting of (in mM) 127 NaCl, 26 NaHCO_3_, 1.6 KCl, 1.24 KH_2_PO_4_, 1.3 MgSO_4_, 2.4 CaCl_2_, and 10 glucose.

### 2.5. In vitro electrophysiology

All recordings were performed at 32−33°C. Whole-cell recordings were performed using ACC layer 2/3 pyramidal cells using a MultiClamp 700B amplifier and a Digidata 1550 digitizer controlled by pCLAMP10.5 software (Molecular Devices). Borosilicate glass pipettes (3−6 MΩ) were filled with a solution containing (in mM) 130 CsMeSO_4_, 10 CsCl, 10 HEPES, 10 phosphocreatine, 4 MgATP, 0.3 NaGTP, and 10 QX-314. Paired whole-cell recordings were performed in two neurons located less than 100 μm apart; one glass pipette contained 200 μM picrotoxin. Liquid junction potentials were not corrected. The signals were gained 5−10-fold, low-pass-filtered at 1 kHz and digitized at 20 kHz.

### 2.6. Drug application

Male mice (6 postnatal weeks) were anesthetized with xylazine (10 mg/kg, i.p.) and pentobarbital (2.5 mg/kg, i.p.). Stainless-steel cannulas (Plastic One) were bilaterally implanted and aimed at the ACC (AP: +1.0 mm, LM: ±0.3 mm, DV: -2.0 mm). The cannulas were chronically secured to the skull using a mixture of acrylic and dental cement. Dummy cannulas (33-gauge) were inserted into each guide cannula to prevent clogging. Mice were allowed to recover for at least 7 days. Before testing, mice received an intra-ACC infusion (0.5 μl per side) of clonazepam (500 nM, Wako). The clonazepam was dissolved in DMSO and was diluted in phosphate-based saline (PBS) containing 0.1 mM SR101 so that the final concentration of DMSO was 0.001%. Infusions were performed slowly over more than 2 min using 33-gauge cannulas. Twenty minutes after infusion, the mice were subjected to the three-chamber test. After the test, the infused regions were confirmed by the track of the cannulas and the fluorescence of SR101.

### 2.7. Histology

Layer 2/3 pyramidal neurons in the ACC were intracellularly perfused with 2 mM biocytin through whole-cell recordings. After the recordings, the slices were fixed for 20 h at 25°C in 0.1 M Na_3_PO_4_, pH 7.4, containing 3% (w/v) formaldehyde and 0.01% (w/v) glutaraldehyde. All subsequent incubations were performed at room temperature and were followed by three 30-min rinses with PBS containing 0.3% (v/v) Triton X-100 (PBS-X). After being blocked with PBS-X containing 1% (v/v) goat serum and 0.12% (w/v) λ-carrageenan (PBS-XCG) for 1 h, the slices were incubated for 48 h with 1/1,000-diluted mouse anti-gephyrin antibody (147021; Synaptic Systems) in PBS-XCG. The slices were further incubated for 24 h with a mixture of 5 μg/mL Alexa Fluor 488-conjugated streptavidin (S11223; Life Technologies) and 10 μg/mL Alexa Fluor 647-conjugated goat antibody to mouse IgG (A21236; Life Technologies) in PBS-XCG. The sections were mounted onto gelatinized glass slides and cover-slipped with 50% (v/v) glycerol and 2.5% (w/v) triethylenediamine (antifading reagent) in 20 mM Tris-HCl (pH 7.6). The slices were observed under a Leica TCS SP8 confocal laser-scanning microscope (Leica Microsystems) with a water-immersion objective lens (25×, 0.95 numerical aperture, Leica) and HyD detectors (Leica). We acquired image stacks at depths ranging from 5 to 25 μm relative to the surface with the pinhole at 1.0 airy disk unit and zoom factor at 1. The image stacks were saved as 8-bit TIFF files. The image size was 4,096 × 4,096 pixels, corresponding to a pixel dimension of 114 nm in both the X and Y directions. Alexa Fluor 488 and Alexa Fluor 647 were excited with 488-nm and 638-nm laser beams and observed through 500–550 and 650–800 nm emission prism windows, respectively. Up to 20 images were acquired at a *Z*-interval of 1 μm per stack and were deconvoluted using Huygens Essential software (version 3.7; Scientific Volume Imaging).

## 3. Results

### 3.1. Poly(I:C)-treated mice are less social

We treated pregnant mice with poly(I:C), a synthetic analogue of double-stranded RNA that mimics a molecular pattern in viral infection, and used the offspring of these mice as the MIA model (Naviaux et al., 2013). We first monitored the exploratory behavior of these mice in a three-chamber maze in which one chamber contained an unfamiliar mouse in the corner box and nothing in the other box (Fig. 1A). Control mice whose mothers had been treated with saline during their embryonic periods preferentially explored the chamber with the unfamiliar mouse, but mice whose mothers had been treated with poly(I:C) (hereafter termed poly(I:C)-treated mice) spent less time in the chamber containing the mouse (Fig. 1B, *P* = 0.016, *D*_19_ = 0.44, Kolmogorov-Smirnov test, *n* = 11 control and 10 poly(I:C) mice). The total time spent exploring both the mouse-containing and vacant chambers did not differ between these two groups (Fig. 1B inset; *P* = 0.87, *t*_19_ = 0.19, Student’s *t*-test), meaning the spatial exploration motivation was intact in poly(I:C)-treated mice.

**Figure 1.**
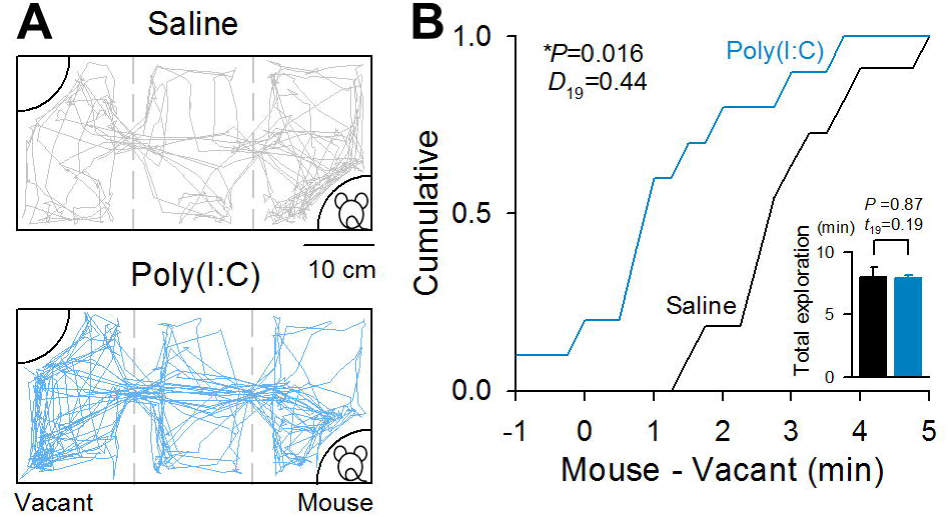
Poly(I:C)-treated mice exhibit low social interaction. **A,** The diagram depicts the exploration tracks of control and poly(I:C)-treated mice during a three-chamber test for a total of 10 min. Mice were treated twice with saline or poly(I:C) on embryonic days 12.5 and 17.5. During the test, an unfamiliar mouse was presented in the corner of the right chamber; the left chamber was vacant. **B,** The graph plots the cumulative distributions of the difference between the times spent in the left and right chambers by 11 saline-treated and 10 poly(I:C)-treated mice. The inset indicates the total time spent exploring the left and right chambers. Kolmogorov-Smirnov test.

### 3.2. Excitatory inputs to ACC pyramidal cells are increased in poly(I:C)-treated mice

Hyperexcitation of synaptic transmission is implied in neuropsychiatric symptoms (Gogolla et al., 2009, Rubenstein and Merzenich, 2003). The ACC is a brain region related to social behavior (Lamm et al., 2011). Thus, we prepared acute ACC slices and recorded spontaneous EPSCs from layer 2/3 pyramidal cells (Fig. 2A). The EPSC frequencies were higher in slices from poly(I:C)-treated mice than in those from control mice (Fig. 2B; *P* = 1.6 × 10^−3^, *t*_17_ = 3.76, Student’s *t*-test; *n* = 9 control and 10 poly(I:C) cells). The EPSC amplitudes were also higher in poly(I:C)-treated mice than in control mice (Fig. 2C, *P* = 1.9 × 10^−2^, *t*_17_ = 2.60, Student’s *t*-test; n = 9 control and 10 poly(I:C) cells).

**Figure 2.**
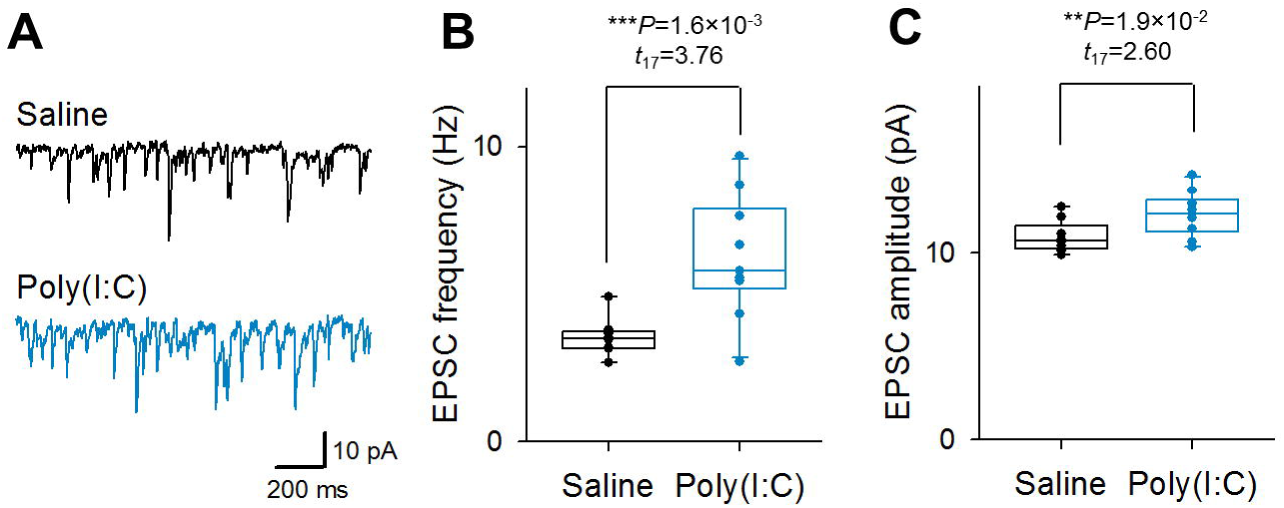
ACC neurons are hyperactive in poly(I:C)-treated mice. **A,** Representative EPSC traces in layer 2/3 pyramidal cells in acute slices prepared from control and poly(I:C)-treated mice. **B,** Frequency of spontaneous EPSCs in layer 2/3 pyramidal cells in acute ACC slices prepared from control and poly(I:C)-treated mice. The data were calculated from 9 neurons from 5 saline-treated mice and 10 neurons from 10 poly(I:C)-treated mice. Student’s *t*-test. **C,** The same as B but for the amplitudes of spontaneous EPSCs.

### 3.3. Excitatory synaptic connectivity in the ACC is intact in poly(I:C)-treated mice

We next focused on excitatory synaptic connections between ACC layer 2/3 pyramidal cells. We conducted double whole-cell recordings from ACC layer 2/3 pyramidal cells. The synaptic connection probability did not differ between control and poly(I:C)-treated mice (Fig. 3A; *P* = 0.40, Fisher’s exact test). The synaptic strength between connected cell pairs did not differ significantly (Fig. 3B; *P* = 0.093, *t*_12_ = 0.09, Student’s *t*-test). These results suggest that excitatory synaptic connectivity in the ACC was not affected in poly(I:C)-treated mice.

**Figure 3.**
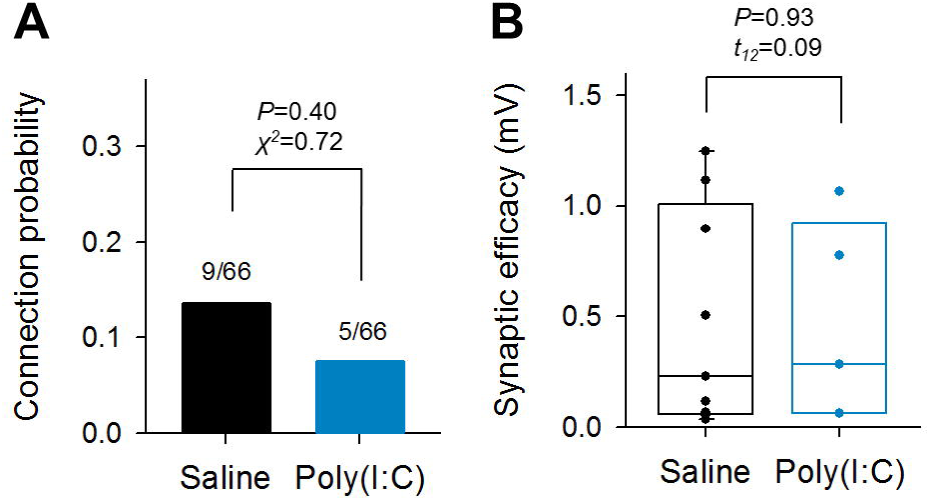
Excitatory connections are not changed in the ACC of poly(I:C)-treated mice. **A,** The probability of an excitatory monosynaptic connection in ACC layer 2/3 pyramidal cells prepared from control and poly(I:C)-treated mice. Chi-squared test. **B,** The synaptic efficacy is the average of evoked monosynaptic EPSPs containing synaptic failure. They were measured from 9 neurons in control mice and 5 neurons in poly(I:C)-treated mice. Student’s *t*-test.

### 3.4. Inhibitory conductance in the ACC is reduced in poly(I:C)-treated mice

We next focused on inhibitory synapses. The immunoreactivity of gephyrin, a protein that anchors inhibitory receptors, was reduced in the somata and dendrites of layer 2/3 pyramidal cells in the ACC of poly(I:C)-treated mice (Fig. 4; Soma: *P* = 5.0 × 10^−3^, *U* = 279; Dendrite: *P* = 0.043, *U* = 262, Mann-Whitney’s *U*-test). We recorded spontaneous EPSCs from ACC layer 2/3 pyramidal cells in which 200 μM picrotoxin, a GABA_A_ receptor channel blocker, was intracellularly injected to reduce local GABAergic influences (Inomata et al., 1988, Nelson et al., 1994). Note that we employed intracellular picrotoxin loading instead of direct recording of inhibitory synaptic transmission because voltage-clamp recording at 0 mV can isolate only peri-somatic inhibitory inputs (Williams and Mitchell, 2008), whereas intracellularly perfused picrotoxin can affect even GABA_A_ receptors located in distal dendrites. In slices from control mice, the picrotoxin-loaded neurons exhibited higher EPSC frequencies than the simultaneously recorded control neurons (Fig. 5A; *P* = 6.6 × 10^−3^, *t*_10_ = 3.42, paired *t*-test; n = 11 pairs). By contrast, in slices from poly(I:C)-treated mice, intracellular perfusion with picrotoxin did not further increase the EPSC frequency (Fig. 5B; *P* = 0.29, *t*_14_ = 1.09, paired *t*-test; *n* = 15 pairs). These results suggest malfunction of GABA_A_ receptors in MIA-model mice.

**Figure 4.**
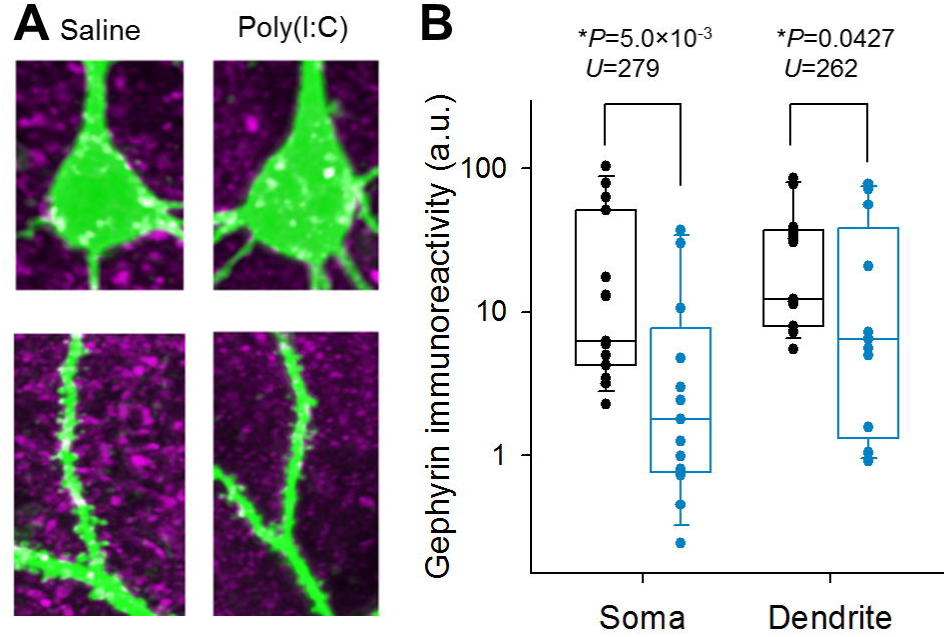
Gephyrin expression is reduced in the ACC of poly(I:C)-treated mice. **A,** Representative confocal image of fluorescent immunoreactivity of gephyrin (*magenta*) in the somata and dendrites of layer 2/3 pyramidal cells in mice whose mothers had been treated with saline or poly(I:C) during the embryonic period. Neurons were visualized with Alexa Fluor 488-conjugated streptavidin (*green*). Biocytin was injected through patch-clamp pipettes into the cell. **B,** Gephyrin immunoreactivity in Alexa-injected neurons was measured in the somata and dendrites of 15 and 13 neurons in 3 control and 3 poly(I:C)-treated mice. Mann-Whitney *U*-test.

**Figure 5.**
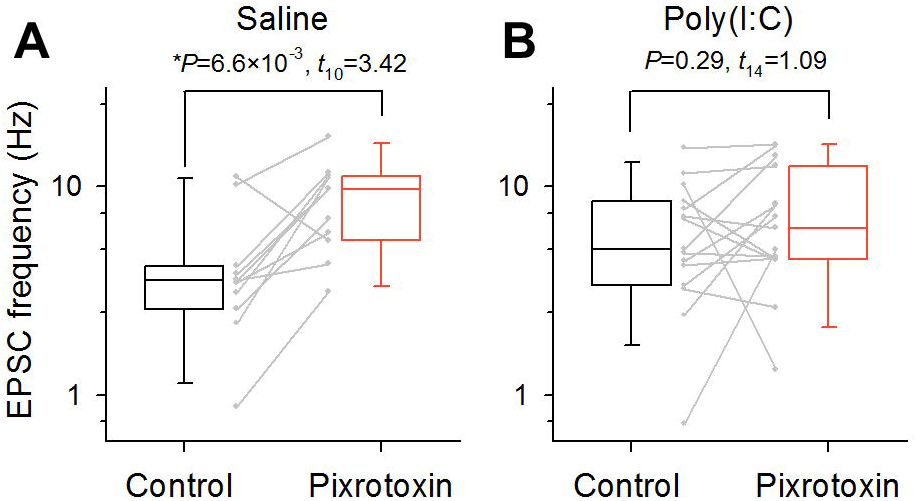
Inhibitory conductance is decreased in poly(I:C)-treated mice. Dual recordings of spontaneous EPSCs from ACC layer 2/3 pyramidal cells in control (**A**) and poly(I:C)-treated (**B**) mice. In each pair, one neuron was intracellularly perfused with 200 μM picrotoxin. The data are box plots of 11 pairs from 10 saline-treated mice and 15 pairs from 10 poly(I:C)-treated mice. Paired *t*-test.

### 3.5. Clonazepam treatment in the ACC recues social behavior

We bilaterally injected clonazepam, a pharmacological enhancer of GABA_A_ receptors that contain α2/3-γ2 subunits (Benke et al., 1996, McKernan et al., 1991), into the ACC of poly(I:C)-treated mice at a dose of 80 pg/site 20 min before the behavioral test (Fig. 6A). The poly(I:C) mice treated with clonazepam injection explored the mouse chamber more preferentially than vehicle-treated poly(I:C) mice (Fig. 6B,C; *P* = 0.016, *D*_20_ = 0.44, *n* = 11 poly(I:C)-treated mice), although the total exploration time was not different between the two groups (Fig. 6B inset; *P* = 0.92, *t*_20_ = 0.10, paired *t*-test). Thus, hyperexcitation of the ACC is responsible for social deficits in MIA-model mice.

**Figure 6.**
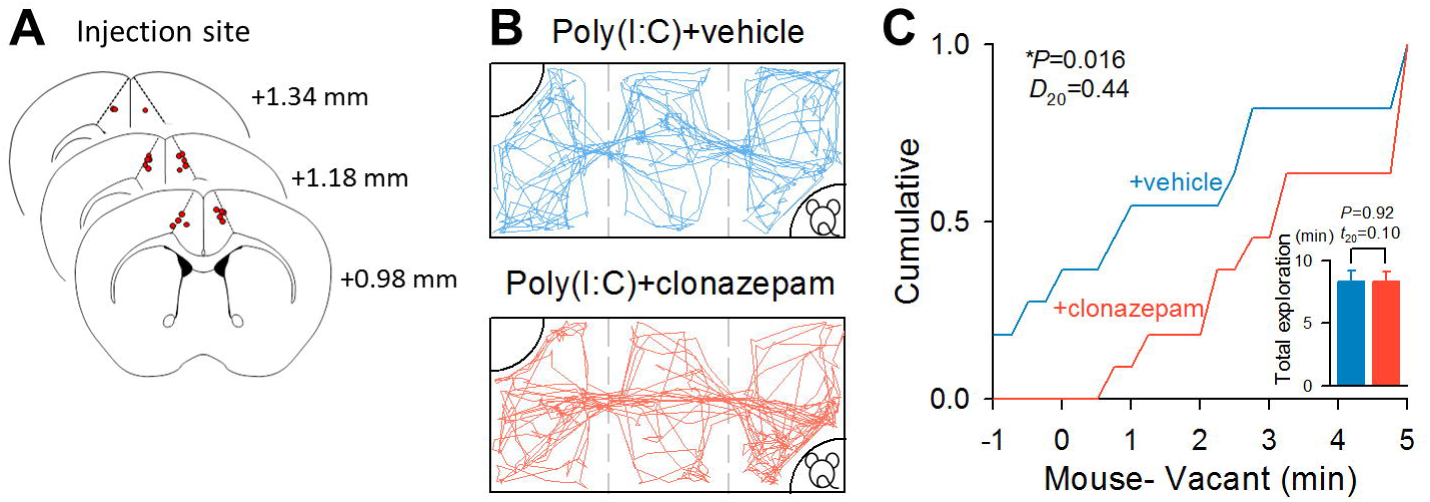
GABA_A_ receptor activation in the ACC recovers social behavior. **A,** *Post hoc* histological inspection reveals successful cannula implantation in the ACC. The target locations of local injection of clonazepam are indicated by the red circles (*n* = 11 mice). The locations were confirmed based on the track of the cannulas and the infusion of 0.1 mM SR101. **B,** The same diagram as in Figure 1, depicting the exploration tracks of control and poly(I:C)-treated mice during a three-chamber test for a total of 10 min, but for 11 poly(I:C)-treated mice administered vehicle or clonazepam in the bilateral ACC 20 min before the behavioral test. **C,** The graph plots the cumulative distributions of the difference between the times spent in the left and right chambers by 11 vehicle-or clonazepam-treated poly(I:C)-treated mice. The inset indicates the total time spent exploring the left and right chambers. Kolmogorov-Smirnov test.

## 4. Discussion

We found that the ACC in MIA-model mice is hyperexcitable due to a reduction in inhibitory inputs and that strengthening the inhibition of the ACC of these mice restores their social deficits. These results suggest that GABAergic malfunction in the ACC underlies social deficits of neuropsychiatric disorders.

We measured the sociability of poly(I:C)-treated mice using the three-chamber behavior test. For these behavioral experiments, we used juvenile mice, which are younger than those generally used in the three-chamber test in previous studies. Our results indicate that the MIA-model mice exhibit social deficits even at an early age.

Accumulating evidence has suggested that neuronal hyperexcitation caused by the net imbalance between excitation and inhibition underlies nervous system disorders in humans, including epilepsy, mental retardation, cognitive impairment, and ASD (Eichler and Meier, 2008, Verret et al., 2012). In poly(I:C)-treated MIA-model mice, cortical parvalbumin-positive interneurons are lacking, and this deficit leads to behavioral abnormalities, including reversing behavior and social deficits (Yim et al., 2017). We found that ACC neurons were physiologically hyperactive due to the hypofunction of GABAergic synapses in poly(I:C)-treated mice. This hyperexcitation of ACC neurons may cause the social deficits of poly(I:C)-treated mice because the pharmacological enhancement of GABAergic transmission in the ACC alone was sufficient to attenuate their social dysfunction. A recent study revealed that inhibition of the primary somatosensory cortex is also sufficient to restore the social interactions (Yim et al., 2017). Social behavior is complex and is unlikely generated in a single brain region. Because brain regions are connected in a complex manner, it is possible that specific inhibition of ACC affects the excitability of other brain regions, such as the primary somatosensory cortex, and vice versa.

Phasic inhibition, one of the forms of inhibition produced by activation of GABA_A_ receptors, works to efficiently modulate excitatory synaptic activity (Hayama et al., 2013, Liu, 2004, Muller et al., 2012). Thus, a lack of phasic inhibition is expected to lead to neuronal hyperexcitation. The phasic inhibition is generated by synaptic GABA_A_ receptors that contain α2/3-γ2 subunits (Farrant and Nusser, 2005). To restore the phasic inhibition, therefore, we used clonazepam, an enhancer of GABA_A_ receptors that contain α2/3-γ2 subunits. Clonazepam is prescribed for epilepsy and anxiety disorders and was also suggested to ameliorate dementia (Busche et al., 2015). However, it has not been applied for human social disorders. A previous work suggested that intraperitoneal injection of clonazepam restored behavioral deficits in a mouse model of autism (Han et al., 2014). These results together with ours suggest that clonazepam may act on inhibitory transmission of the ACC and ameliorate sociability.

## 6. Acknowledgments

We thank Dr. Chiaki Kobayashi and Dr. Tetsuya Sakaguchi for their comments on the experiments and the analyses. This work was partly supported by Grants-in-Aid for Scientific Research (18H05114 to YI), the Human Frontier Science Program (RGP0019/2016 to YI), and MEXT/JSPS (16H04663, 17K19451, 15H01430, 18H04743 to HH) and by Brain/MINDS from AMED (JP18dm0207064 to HH). This work was conducted partly as a program at the International Research Center for Neurointelligence (WPI-IRCN) of The University of Tokyo Institutes for Advanced Study at The University of Tokyo.

